# Stress drives the hippocampus to prioritize statistical prediction over episodic encoding

**DOI:** 10.1101/2025.10.25.683838

**Authors:** Irene Zhou, Zihan Bai, Yuye Huang, Elaine G. Wijaya, Brynn E. Sherman, Nicholas B. Turk-Browne, Elizabeth V. Goldfarb

**Affiliations:** Department of Psychology, Yale University; Department of Psychology, New York University; Department of Psychological & Brain Sciences, Johns Hopkins University; School of Medicine, UC Riverside; Department of Psychology, The Ohio State University; Department of Psychology, Wu Tsai Institute, Yale University; Department of Psychiatry, Wu Tsai Institute, Yale University

## Abstract

The hippocampus plays a critical role in encoding individual experiences into episodic memory, while also detecting patterns shared across these experiences that allow us to anticipate future events via statistical learning. Acute stress is known to impact the hippocampus but may have opposite effects on these competing functions. That is, although stress impairs episodic encoding, it may instead enhance statistical learning. Using functional magnetic resonance imaging, we tested how stress influences the hippocampal pathways and subfields that support episodic encoding and statistical learning while participants performed a task that engaged both processes. Across several analyses, stress biased hippocampal processing in favour of statistical learning, suppressing pattern separation between events in the dentate gyrus and enhancing the representation of statistically predictive features. Furthermore, stress drove the hippocampal monosynaptic pathway to support statistical learning rather than episodic encoding during predictive situations. Together, these data suggest that acute stress elicits targeted, adaptive changes in hippocampal pathways which may facilitate predicting and responding to upcoming events.

## 1 Introduction

Our lives contain many experiences that are unique but unfold in a reliable way. For example, a tree with a double-blaze marking on a hiking trail is often followed by a sharp turn in the trail. These reliable patterns can be detected automatically through statistical learning, a process which gradually extracts features that are shared across multiple experiences. Learning these regularities allows us to detect predictive situations and anticipate future events (looking for a hidden turn in advance). At the same time, unique details of individual experiences — such as a friend you encounter unexpectedly — are encoded rapidly and distinctively in episodic memory.

Despite episodic encoding and statistical learning operating on different timescales (fast versus slow, respectively) and prioritizing different features (unique versus shared, respectively), the hippocampus is thought to support both functions, albeit along distinct pathways. Neural network models suggest that statistical learning is supported by the monosynaptic pathway (MSP), which is a direct, recurrent connection between entorhinal cortex (EC) and hippocampal subfield CA1 that produces overlapping representations of related experiences over time. In contrast, episodic encoding is supported by the trisynaptic pathway (TSP), which connects EC to CA1 via intermediary connections through dentate gyrus (DG) and CA3 [1]. Functional neuroimaging supports this dual function, with hippocampal patterns supporting both predictive representations [2, 3] and unique event signatures [4, 5]. Although these learning processes are supported by different neural pathways, their shared dependence on CA1 as well as entorhinal inputs and outputs suggests that they may compete for shared neural resources that determine the representations and output of the hippocampus. Indeed, there are behavioural and neural trade-offs between making predictions and forming individual episodic memories [3].

The role of stress in arbitrating between these hippocampal functions is unknown. The hippocampus has long been known to be sensitive to acute stress [6, 7, 8]. Acute stress reduces plasticity [9] and elicits rapid dendritic atrophy [10, 11] in rodent hippocampal neurons. In humans, acute stress reduces haemodynamic activity in the hippocampus and medial temporal lobe cortex [12, 13] and alters hippocampal connectivity [14, 15, 16]. Such actions have been used to explain deleterious stress effects on episodic encoding [17, 18], particularly for stimuli that are emotionally neutral and unrelated to the stressor [19]. However, recent studies suggest that statistical learning behaviour can be enhanced — rather than impaired — by stress [20, 21], raising the possibility that stress exerts differential effects across hippocampal pathways.

A close examination of stress effects on subfields in the hippocampus suggests how stress might have opposite effects on episodic encoding and statistical learning. Across species, acute stress impairs adult hippocampal neurogenesis [22, 23, 24], which contributes to episodic encoding by supporting pattern separation — the formation of sparse, orthogonalized memory traces in DG to minimize interference [25, 26, 27]. Furthermore, acute stress and stress-related elevations in glucocorticoids suppress long-term potentiation in DG [28] and CA3 [29, 30]. Notably, patterns differ for CA1, which has a lower density of glucocorticoid receptors [8, 6]. Indeed, acute stress and glucocorticoids can spare or even enhance long-term potentiation in CA1 [29, 31], facilitating neuronal firing in CA1 [32, 33] but not in DG [34]. Together, these findings suggest that acute stress may impair subfields and connections involved in TSP, while sparing or enhancing those associated with MSP. However, the differential effects of acute stress on these hippocampal pathways in humans has not been tested.

The present study examines how acute stress affects hippocampal processes supporting episodic memory and statistical learning by combining functional magnetic resonance imaging (fMRI) with a task that assesses both functions simultaneously. We predict that stress impairs episodic encoding associated with TSP while sparing competing statistical learning processes supported by MSP. After either acute stress or a matched control procedure, participants viewed a stream of unique images structured into pairs of predictive and predictable scene categories [3]. The next day (to limit acute stress effects on retrieval [18, 35]), participants completed behavioural tests assessing episodic memory (recognizing unique images) and retention of statistical learning (familiarity with pairs). We find that stress alters processing of predictive items across activity patterns within hippocampal subfields, as well as momentary connectivity along hippocampal pathways, with stress consistently prioritizing processing of statistically predictive information at the expense of episodic detail.

## 2 Methods

### 2.1 Participants

Participants were recruited from the New Haven community and pseudorandomly assigned to the stress or control groups to ensure balance in age, sex, and current subjective stress (Perceived Stress Scale [PSS] score). All participants were fluent in English, had BMI between 18 and 35, and had normal or corrected-to-normal vision. To reduce factors that could influence stress-induced cortisol responses, participants were excluded if they were currently using psychiatric, *β*-blocker, or corticosteroid medications, or met criteria for any substance-use disorder. Female participants were not peri- or post-menopausal, pregnant, or lactating, and had not had a hysterectomy.

A total of 96 participants completed both experimental sessions (N = 49 stress [41% male, age 18 – 40, median age 23] N = 47 control [40% male, age 18 – 45, median age 24]; sample size based on a power analysis from a prior study using a similar task [*d* = 0.42, 80% power; [3]]). Groups did not significantly differ in age (*t*(94) = 1.40, *p* = 0.166), sex (*χ*^2^(1, 96) = 0.0, *p* = 1.0), or PSS (*t*(94) = 0.69, *p* = 0.490). Of these participants, *N* = 4 stress and *N* = 4 control were excluded from fMRI analyses due to excess head motion, failure to complete the pre-learning and/or post-learning run, or inattentive responding (<60% of trials with response during stimulus presentation). One additional participant in the control group was excluded from fMRI analyses involving the learning runs because of a technical issue with fMRI data acquisition.

### 2.2 Study procedure

The experimental procedure is shown in Figure 1A. Each participant completed two sessions on consecutive days. Before each session, they acclimated to the environment for at least 10 min (during which they completed the PSS [36]), allowing cortisol levels to stabilize [37].

**Figure 1:**
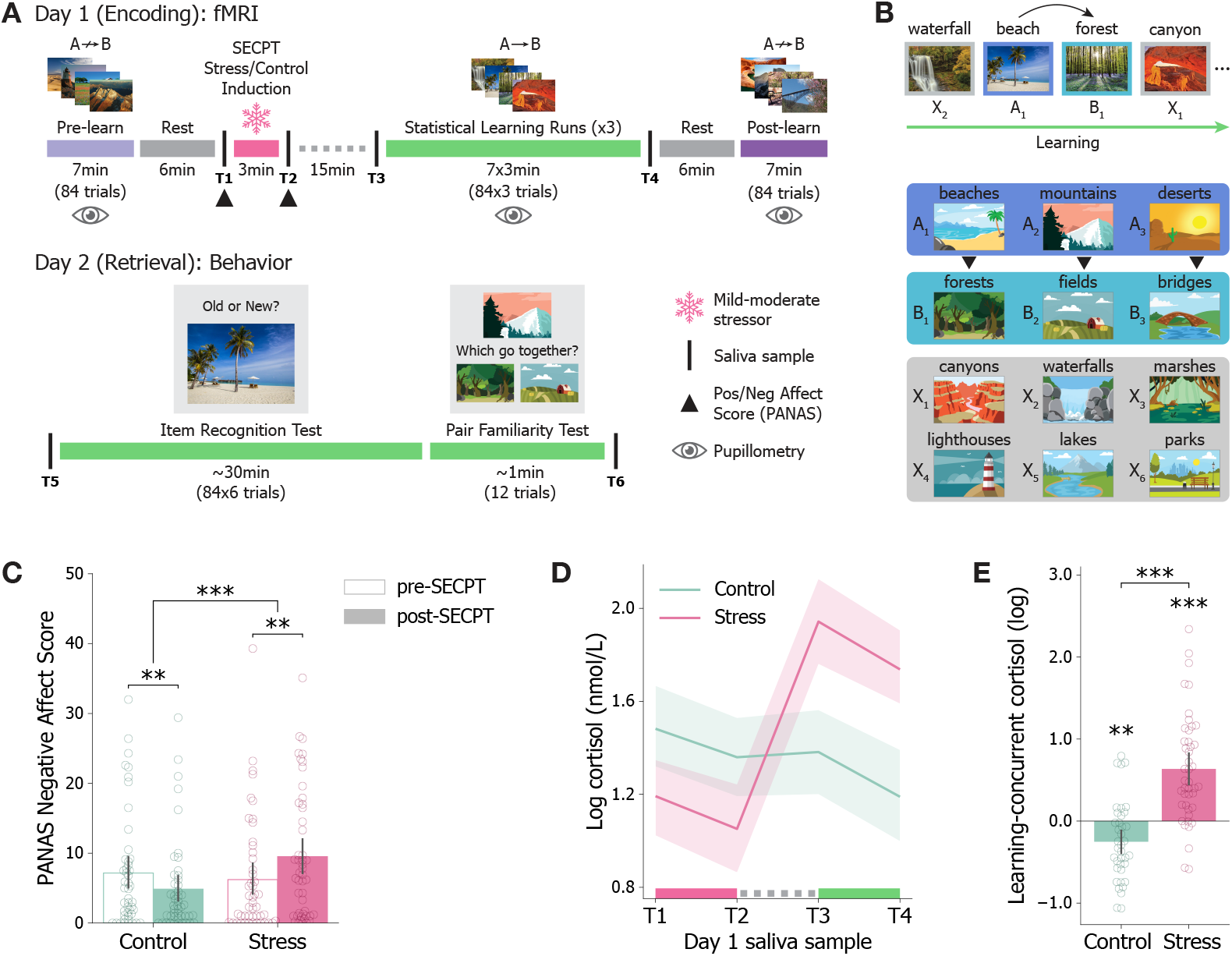
Task design and stress validation. **(A)** Experiment timeline. **(B)** Example trial sequence during statistical learning runs (top). A category images were always followed by paired B category images. Example scene categories from A (predictive), B (predictable), and X (baseline) conditions (bottom). **(C)** Subjective negative affect ratings before and after the SECPT (pink) or matched control procedure (green). **(D)** Salivary cortisol throughout Day 1: T1 (pre-SECPT); T2 (post-SECPT); T3 (pre-learning); T4 (post-learning). **(E)** Mean salivary cortisol across learning (T3 and T4) relative to baseline (T1). Dots = individual participants; error bars = bootstrapped 95% CI across participants. \*\*\**p <* 0.001, \*\**p <* 0.01

On Day 1, participants completed an encoding session during fMRI. During this session, participants were exposed to acute stress (*N* = 49) or a control manipulation (warm water bath; *N* = 47) [38]. After a 15-min delay [20], participants encoded sequences of unique scenes containing temporal regularities across three statistical learning runs. Before the stress induction, participants first completed a pre-learning scene task containing no temporal regularities, followed by a 6-minute period of awake rest. After learning, participants again underwent a period of awake rest, followed by a post-learning run without regularities. Pupillometry data were collected during all scene-viewing runs as a non-invasive, online behavioural measure of sensitivity to statistical regularities.

On Day 2, participants completed an item recognition test (assessing episodic memory) and a pair familiarity test (retention of statistical learning). All experimental sessions occurred between 12:00 PM and 6:00 PM to control for circadian fluctuations in cortisol [7].

#### 2.2.1 Stress induction and measurement

Participants in the stress group underwent the Socially Evaluated Cold Pressor Test (SECPT; [39]). Participants submerged their right arm in a bucket of ice water for 3 min (mean temperature = 2.79° C, SD = 1.02) under social evaluation, which involved being monitored by an experimenter with neutral affect wearing a laboratory coat while being recorded with a front-facing camera. Participants were told that the video recording would be used to evaluate their facial expressions and that they should look into the camera without speaking. Participants were not told how much time remained in the SECPT and were required to keep their arms submerged for the full 3 min. Participants in the control condition underwent a matched induction with warm water (mean temperature = 36.28° C, SD = 4.21) and no social evaluation (neither monitoring nor video recording).

Immediately before and after the stress/control procedure, participants completed the Positive and Negative Affect Schedule (PANAS) to assess changes in subjective affect. Participants also provided salivary samples throughout both visits by placing a Sarstedt salivette under their tongue for 2 mins (Figure 1A). Samples were stored in a −20° C freezer in sterile tubes, then shipped frozen to the laboratory of Dr. Andrea Gierens at the University of Trier for analysis of cortisol via time-resolved fluorescence immunoassay [40]. Because of insufficient sample, 23 of 384 samples from Day 1 (from 15 participants, 6% of total) and 10 of 192 samples from Day 2 (from 9 participants, 5% of total) could not be assayed. A further 3 samples from Day 1, and 2 samples from Day 2 (all from the same participant) were excluded from analyses due to outlier cortisol levels (>4 SD above mean across groups).

#### 2.2.2 Encoding task

The Day 1 encoding task was based on [3] and [20], and was conducted during fMRI. In each trial of the scene-viewing runs (pre-learning, learning, and post-learning), participants viewed a photograph of a scene (1 s) and judged whether the scene contained manmade objects using an MR-compatible button box (response mappings counterbalanced). Trials were separated by 2, 3.5, or 5 s intertrial fixations to jitter onsets for deconvolving event-related fMRI activity.

Each scene presented during encoding was trial-unique but drawn from one of 12 outdoor scene categories (beaches, bridges, canyons, deserts, fields, forests, lakes, lighthouses, marshes, mountains, parks, and waterfalls; Figure 1B). Across the three statistical learning runs, scene categories were paired such that some categories (predictive; A, 3/12 categories) reliably preceded other categories (predictable; B, 3/12 categories) with a transition probability of 1.0 (Figure 1B). Scenes from additional control (X, 6/12 categories) categories (neither predictive nor predictable) were interspersed randomly throughout the sequence (but never between an A-B pair), with assignment of categories to A/B/X conditions randomized per participant. The order of the image sequence within each statistical learning run was randomized with the following three constraints: category pairs and pairs of category pairs could not repeat back to back (i.e., no *A*_1_*B*_1_*A*_1_*B*_1_ or *A*_1_*B*_1_*A*_2_*B*_2_*A*_1_*B*_1_*A*_2_*B*_2_); repetitions of each category were spread equally across quartiles to minimize differences in study-test lag between categories; and the overall transition probability between “yes” and “no” responses on the manmade cover task was forced to be statistically indistinguishable from 0.5.

During pre- and post-learning runs, scenes from each category were presented individually in a random order, with no category pairs. These runs were used to obtain neural templates of category-level representations while avoiding active learning and temporal autocorrelation between A-B pairs. Participants were not informed of statistical regularities. Seven scenes from each scene category appeared in each scene-viewing run (84 scenes each in pre- and post-learning; 252 across learning runs).

#### 2.2.3 Item recognition test

Episodic memory for each trial-unique scene encountered during learning was measured via an old/new recognition memory test on Day 2. In each trial, participants were asked to judge whether the presented scene was “old” (i.e. encountered on Day 1) or “new” (i.e. not seen on Day 1; Figure 1A, bottom left). After making an old/new response, participants rated their confidence (“very unsure”, “unsure”, “sure”, “very sure”). All 252 scene images from the learning runs plus 252 foils (21 novel images from each category) were shown in a random order. Trials terminated following participant response or 6 s (whichever came earlier) and were followed by a 1 s intertrial fixation.

#### 2.2.4 Pair familiarity test

Retention of statistically predictable scene categories was measured via a pair familiarity test. On each trial, a cartoon sketch of an A or B scene category (probe) was presented at the top of the screen (cartoons were used to avoid presenting a familiar or novel exemplar). Cartoons depicting two other scene categories — the paired category (match) and a category from the same condition in another pair (foil) — were presented at the bottom of the screen (e.g., if the probe is an A category, the match and foil are both B categories). Participants were asked to choose which category “goes with” the probe (Figure 1A, bottom right). Each probe/match/foil combination was tested once, resulting in four trials per category pair and 12 trials total. The positions of the match and foil categories were counterbalanced across trials. Each trial terminated following participant response or 6 s (whichever came earlier) followed by a 1 s intertrial fixation.

### 2.3 Data acquisition

#### 2.3.1 Pupil acquisition parameters

Pupillometry data were collected in a subset of participants (N = 33 stress, 28 control) because of technical constraints (primarily, the need for vision-corrective eyewear compatible with eye-tracking and for a clean line of sight to eye-tracking through the head coil). Pupil size was recorded continuously from one eye during scene-viewing runs on Day 1 using an Eyelink 1000 infrared eye-tracker. Pupil size was recorded as the area of a model centroid in arbitrary units (AUs) specified by the eye-tracker’s firmware. Nine-point calibration and validation were performed each time the participant entered the scanner (immediately before the pre-learning run, and immediately before the first learning run) to ensure proper measurement of eye position.

#### 2.3.2 fMRI acquisition parameters

Data were acquired at Yale University on a Siemens Prisma 3T MRI scanner with 64-channel head coil at the Brain Imaging Center in the Faculty of Arts and Sciences (N with exclusions = 21 stress, 21 control) and at BrainWorks in the Center for Neurocognition and Behavior at the Wu Tsai Institute (N with exclusions = 24 stress, 22 control). Participants did not differ significantly across the two sites in age (t(86)=-1.26, p=0.211) or condition assignment (*χ*^2^(1, 88)=0.0, p=1.0). Across all fMRI analyses, there were no significant effects or interactions with scanner site.

Functional images were acquired using an echo-planar imaging (EPI) sequence with the following parameters: repetition time (TR) = 1,500 ms; echo time (TE) = 32 ms; 90 axial slices; voxel size = 1.5 mm isotropic; flip angle = 64°; multiband factor = 6. Anatomical scans included a T1-weighted magnetization prepared rapid gradient echo (MPRAGE) sequence (TR = 2,400 ms; TE = 2.4 ms; voxel size = 1 *×* 1 *×* 1 mm; 208 sagittal slices; flip angle = 8°) and a T2-weighted turbo spin echo (TSE) sequence (TR = 11,170 ms; TE = 93 ms; 54 coronal slices; voxel size = 0.44 *×* 0.44 *×* 1.5 mm; distance factor = 20%; flip angle = 150°).

### 2.4 Data analysis

#### 2.4.1 Quantifying cortisol responses

Salivary cortisol levels were examined to assess physiological stress response following the SECPT. Due to skew, cortisol values were first normalized using a natural-log transform [41]. Differences in cortisol trajectory between groups were assessed using a linear mixed effects model (R 4.5.1, nlme package [42]) predicting log-transformed cortisol as a function of group, timepoint, and their interaction, with participant as a random effect. We further summarized cortisol levels during learning (“learning-concurrent cortisol”; mean(T3, T4) - T1; Figure 1A). Participants with successful assays across T1, T3, and T4 were included in this analysis (N = 38 control, 43 stress).

#### 2.4.2 Behavioural analyses

Performance on the item recognition test was summarized as hit rate, false alarm rate, and A’, a nonparametric measure of sensitivity [43]. Metrics were calculated separately for each scene condition (A, B, X). For the pair familiarity test, participant-level performance was summarized as the mean response accuracy overall, and pair-level performance was summarized as mean accuracy across the four trials for a given pair (separately per participant).

#### 2.4.3 Pupil preprocessing

Pupil data were preprocessed in R (4.5.1) using the gazer package [44] following recommendations from previously established protocols [45]. A subset of 599 trials (2.92% of all trials) were excluded from analysis because they contained less than 50% usable raw datapoints during scene presentation; an additional 144 trials (0.70% of all trials) were excluded because of a lack of participant button press response to the cover task. Blinks and artifacts were detected using a velocity-based algorithm, and data were linearly interpolated during, 20 ms before, and 20 ms after blinks. Data were baseline-corrected at the trial level by subtracting the mean pupil size 0-50 ms after stimulus onset. Pupil size timeseries were then downsampled to 100 Hz.

#### 2.4.4 Pupil analysis

Pupil size was used to examine differences in the online processing of different scene conditions during statistical learning runs. After preprocessing, the mean pupil size trace per condition from the pre-learning run was subtracted from that of the learning runs within participants (traces visualized in 2A). This ensured that condition-level differences in pupil size reflected learning. Data were then averaged per trial (across the 1 s stimulus presentation window) then per condition for each participant. Mean pupil sizes were compared between conditions using bootstrapping within-participant (see section *Bootstrap Statistical Testing*). Group-level differences were assessed using LME to model mean pupil size as a function of group, condition, and their interaction, with participant as a random effect.

#### 2.4.5 fMRI preprocessing

fMRI data were preprocessed using FSL 6.0.3. A total of 6 runs from 3 participants (1 control, 2 stress) were above the a priori motion threshold of *<*1.5 mm absolute mean frame-to-frame displacement as computed by FSL MCFLIRT [46]. These participants were excluded from all fMRI analyses. Data were skull-stripped, pre-whitened, and high-pass filtered at 0.01 Hz to remove low-frequency signal drift. FSL FEAT was then applied to regress out 6 linear estimated motion parameters and white matter timeseries plus their temporal derivatives as well as stick function regressors for nonlinear motion outliers. Residuals were aligned to the participant’s first functional scan. For all analyses, trial-evoked responses were normalized and time-shifted by 3 TRs (4.5 s) relative to task timing to account for haemodynamic lag. Trials with no participant response were excluded from all trial-level imaging analyses.

#### 2.4.6 fMRI region of interest definitions

To examine neural representations and connectivity along hippocampal pathways, we segmented entorhinal cortex (EC) and hippocampal subfields CA1, CA2/3, and DG using the automatic segmentation of hippocampal subfields (ASHS) package [47] with the Princeton Young Adult 3T ASHS Atlas [48] from each participant’s T1 and T2-weighted anatomical scans. As we did not have lateralized predictions, region of interest (ROI) masks were combined across hemispheres. These participant-specific ROIs were then aligned into that participant’s functional space. For all scene-viewing runs, preprocessed data were masked to include only voxels within an ROI, then z-scored across time for subsequent analyses.

#### 2.4.7 Predictive category decoding analysis

A multivoxel pattern classification approach was used to determine whether statistical learning led to neural evidence for predicted categories [3]. After preprocessing and extracting data from ROIs, voxel activity patterns from timepoints corresponding to stimulus presentation of B images were extracted from each participant’s pre-learning run as a training set for a classifier. For each participant, binary linear support vector machines (SVMs) were trained to distinguish B categories (e.g., *B*_1_ = forests, *B*_2_ = fields, and *B*_3_ = bridges) using data and labels from the pre-learning run. As each participant encountered 3 distinct A and B categories over the course of the experiment, three different binary classifiers were trained to distinguish between pairs of B categories (*B*_1_ vs. *B*_2_, *B*_1_ vs. *B*_3_, *B*_2_ vs. *B*_3_). SVMs were defined using the SVC function in scikit-learn with a penalty parameter of 1.00.

These classifiers were then tested on neural data while participants viewed the corresponding A images in the post-learning run (e.g., a classifier trained to distinguish *B*_1_ vs. *B*_2_ in pre-learning was tested on *A*_1_ [e.g., beach] vs. *A*_2_ [mountain] trials in post-learning). The post-learning run was used to avoid temporal autocorrelation between paired A and B images in the learning runs (A was always followed by B); the training and testing sets were thus drawn from runs with randomized temporal structure. Classifier accuracy was defined as the percent of correct predictions (e.g., a classifier made a correct prediction if an *A*_1_ scene was classified as *B*_1_ over *B*_2_). Accuracy was averaged over all three classifiers, resulting in one mean accuracy score per participant. To assess reliability at the group level, performance was compared to an assumed chance level of 0.5 (given the binary nature of the classifier) across participants using bootstrapping.

To verify our results without assuming chance accuracy, we performed randomization tests in which we computed an empirical null distribution of classification accuracy values for each participant. Null distributions were generated from 9,999 iterations of shuffling A category labels in each participant’s post-learning run before extracting binary testing sets to score each classifier. Scores were then averaged within participant to produce a mean (null) score for each iteration. We then converted each participant’s true classification accuracy to a z-score relative to their own null distribution, and tested reliability at the group level by comparing z-scored accuracies against 0 using bootstrapping.

Before training and testing each classifier, feature selection was performed by selecting a subset of voxels using the SelectKPercent function in Python’s scikit-learn module. The percentage of voxels selected from each ROI was consistent across all participants (CA1: 12.5%; CA2/3: 20%; DG: 12.5% of voxels). The best value for this percentage parameter was determined across all participants by using nested leave-one-out cross-validation within the training data only.

#### 2.4.8 Category representation similarity analysis

A multivoxel pattern similarity approach was used to assess pattern separation during learning (see Fig. 3B). After preprocessing and extracting data from ROIs, voxel activity patterns from timepoints corresponding to stimulus presentation were extracted from each participant’s learning runs. For each trial, *within-category* pattern similarity was computed as the mean Pearson correlation between the activity pattern for that trial (e.g., a beach) and all other trials from the same scene category (all other beaches) during that learning run. *Between-category* pattern similarity was computed as the mean Pearson correlation between the activity pattern for that trial (e.g., a beach) and all other trials from different categories within the same condition (that is, all non-beach images from other A categories) during that learning run. After normalizing mean correlations using a Fisher r-to-z transform, between-category pattern similarity was subtracted from within-category pattern similarity, and the resulting values were averaged within participant and condition. Values less than or equal to 0 reflect pattern separation, indicating that scenes within the same category were represented more distinctly than scenes from different categories. Group-level evidence for pattern separation was assessed by comparing against a null value of 0 across participants using bootstrapping. Within-group differences between conditions were assessed using paired-samples bootstrapping, and within-condition comparisons across groups were assessed using between-groups bootstrapping.

#### 2.4.9 Category evidence analysis

Multivoxel pattern similarity was also used to quantify category-level evidence in each learning trial (see Fig. 3C), or the mean Pearson correlation between the activity pattern for that trial and all trials with the same scene category from the pre-learning run (e.g., a beach from learning compared to every beach from pre-learning). Higher values indicated greater similarity between a particular item and the category overall, or more of a category-level representation. Resulting trialwise category evidence values were then normalized using a Fisher r-to-z transform then averaged within participant and condition. As in *Category representation similarity analysis*, comparisons against 0, within-group differences between conditions, and within-condition comparisons across groups were assessed using bootstrapping.

#### 2.4.10 Momentary cofluctuation analyses

In addition to examining patterns within subfields during learning, we also calculated functional coupling between subfields within hippocampal pathways. Given their overlapping role in both MSP and TSP, we focused on coupling between CA1 and EC. After preprocessing, we used a GLM to fit stimulus-evoked responses with a canonical double-gamma haemodynamic response function (plus temporal derivative) for each trial. We then extracted the residuals of this model (variance not explained by univariate responses) from the voxels in each subfield and calculated the mean time-series across voxels. Using an edge timeseries approach [49, 50], we quantified momentary cofluctuation for pairs of subfields by z-scoring their residual timeseries and multiplying them on a timepoint-by-timepoint basis. Averaging the resulting edge timeseries across time (more specifically, summing and dividing by the total number of timepoints minus 1) yields the Pearson correlation (i.e., background connectivity; see [51, 52, 53]) between subfields. Thus, timepoints with large positive products contribute to more positive overall connectivity, whereas those with large negative products contribute to more negative overall connectivity.

We then determined how connectivity between subfields during predictive images related to episodic encoding and statistical learning. For episodic encoding, we compared momentary average cofluctuation values between subsequently remembered and forgotten A images (that is, correctly identified as “old” and incorrectly identified as “new”, respectively, in the item recognition test on day 2). For statistical learning, we related the average momentary cofluctuation for each predictive A category to behavioural familiarity for the corresponding A-B pair. To facilitate comparison with the episodic memory approach, we repeated the statistical learning analysis after binarizing performance on the pair familiarity test, labelling pairs with above-chance (>0.5) accuracy as “learned” and pairs with below-chance (<0.5) accuracy as “not learned”, and omitting pairs with at-chance (=0.5) accuracy. For the continuous version of this analysis, see Supplemental Figure S4.

#### 2.4.11 Bootstrap statistical testing

Unless otherwise stated, statistical tests were conducted using a non-parametric bootstrap resampling method [54] with 99,999 iterations. For one-sample testing or paired-sample comparisons, participants were resampled with replacement and the group mean computed for each iteration. For between-participants comparisons, participants were resampled with replacement within groups, and the difference between group means was computed for each iteration. For comparisons against chance, bootstrapped p-values were calculated as the proportion of sampling iterations yielding a group mean less than or equal to chance. For two-tailed comparisons (e.g., between groups or conditions), the p-value was calculated as the proportion of sampling iterations yielding a difference in the opposite direction of the true difference, multiplied by 2. For correlations, participants were resampled with replacement and the Pearson correlation between the two variables of interest was recalculated for each iteration. The resulting correlation coefficient (*r*) from each iteration was used to populate a sampling distribution. P-values were calculated as the proportion of sampling iterations in which the sampled correlation was the opposite sign from the true correlation, multiplied by 2 for two-tailed significance. In all cases, 95% confidence intervals were defined as the 2.5th and 97.5th percentiles of the sampling distribution.

## 3 Results

### 3.1 Stress manipulation evoked changes in subjective affect and cortisol

We first evaluated the efficacy of the stress induction by examining changes in subjective affect (PANAS scores; Figure 1C). Participants in the stress group felt significantly more negative after the SECPT (*p* = 0.009, 95% CI: [0.88, 5.63]), whereas participants in the control group reported feeling significantly *less* negative affect (*p* = 0.001, 95% CI: [−4.06, −0.80]). Change in negative affect was significantly higher in the stress group than in the control group (*p <* 0.001, 95% CI: [2.75, 8.48]).

Given the importance of glucocorticoids for stress effects on hippocampal function, we next assessed whether the stress induction increased salivary cortisol levels during learning. Based on previous work [20], we timed the onset of learning relative to SECPT with the expectation that cortisol would be elevated from the beginning to the end of learning. As anticipated, groups showed significantly different cortisol trajectories throughout Day 1 (Figure 1D; main effect of group: *F* (1, 78) = 1.61, *p* = 0.021; main effect of timepoint: *F* (1, 238) = 14.91, *p <* 0.001; group × timepoint: *F* (1, 238) = 74.56, *p <* 0.001). In the stress group, cortisol was significantly elevated during learning (average of pre- and post-learning minus baseline; *p <* 0.001, 95% CI: [0.45, 0.85]), whereas cortisol was significantly reduced during learning in the control group (*p* = 0.002, 95% CI: [−0.40, −0.10]; stress vs control: *p <* 0.001, 95% CI: [0.65, 1.14]). These stress effects were specific to learning, as we did not observe any differences in cortisol between groups on the following day during the retrieval session (main effect of group: *F* (1, 84) = 0.25, *p* = 0.620; main effect of timepoint: *F* (1, 84) = 2.04, *p* = 0.157; group *×* timepoint: *F* (1, 84) = 1.14, *p* = 0.288).

### 3.2 Both groups showed pupillary evidence of statistical learning

We next examined whether participants were sensitive to the regularities in the scene sequence during statistical learning. If participants learned the regularities, they should show differential responses to the A, B, and X scene conditions that were otherwise matched. We anticipated that these differences would be reflected in pupil responses. Indeed, there was a significant difference in mean pupil size between A and B, with the control group exhibiting more constriction (smaller pupil) during B (A vs. B: *p* = 0.018, 95% CI: [2.00, 20.28]; A vs. X: *p* = 0.936, 95% CI: [−12.35, 13.49]; B vs. X: *p* = 0.021, 95% CI: [−19.68, −1.62]) and the stress group exhibiting more constriction during A (A vs. B: *p* = 0.008, 95% CI = [−31.36, −4.32]; A vs. X: *p* = 0.018, 95% CI: [−21.28, −1.85]; B vs. X: *p* = 0.273, 95% CI: [−4.74, 17.55]). Although stress flipped the direction of these effects (group × condition: *F* (2, 118) = 6.12, *p* = 0.003), the observation that participants in both groups showed differential pupillary responses to predictive and predictable images suggests sensitivity to learned regularities.

### 3.3 Both groups showed neural evidence of prediction in CA2/3 after statistical learning

We expected that statistical learning would facilitate prediction of upcoming B categories while viewing paired A categories. Based on previous work [3], we anticipated that this learned prediction would be evident in hippocampus; within the hippocampus, such effects have been observed reliably in the CA2/3 subfield [2, 55, 56]. We thus tested whether expected B categories could be reliably decoded from patterns of activity in hippocampal subfields during A images *after* learning (using a classifier trained to separate B categories *before* learning; approach shown in Figure 2B).

**Figure 2:**
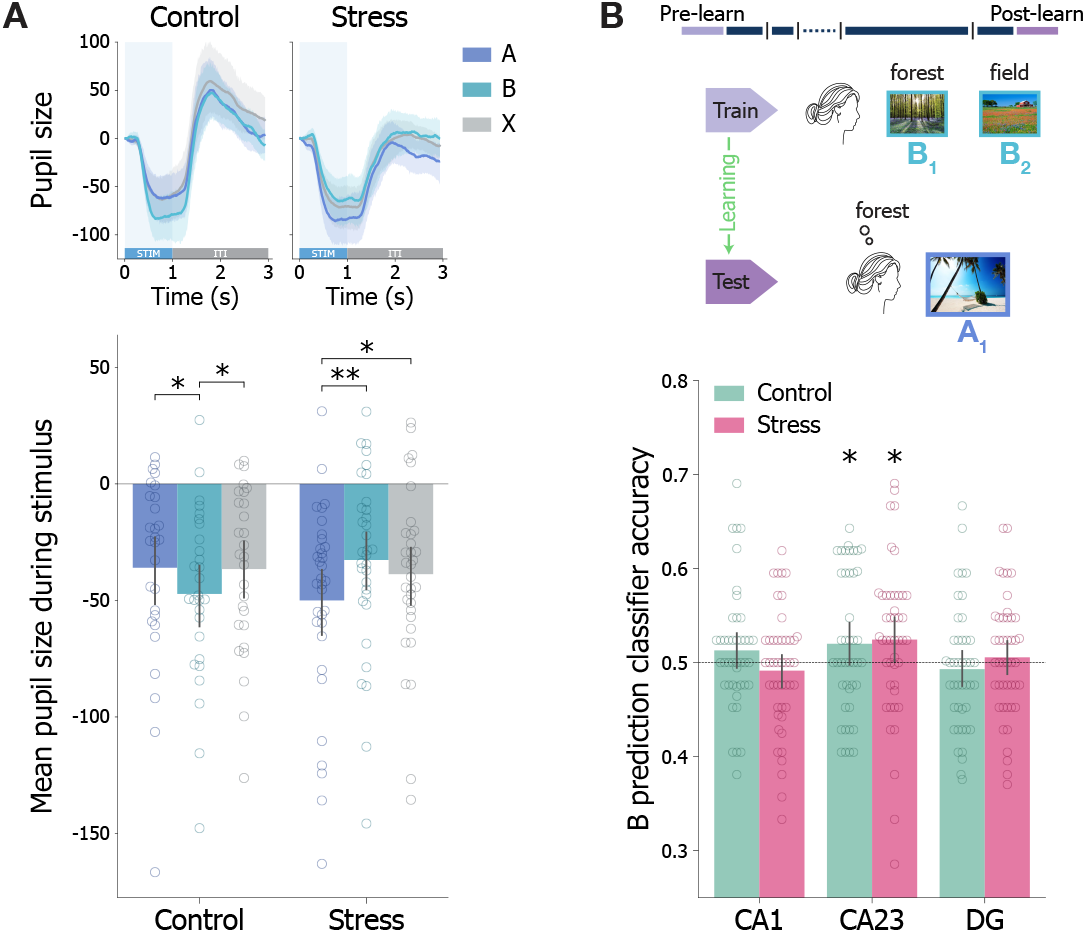
Online and offline evidence of statistical learning. **(A)** Pupil size for A, B, and X scenes during learning (top; blue shading = stimulus presentation, grey = ITI). Mean pupil size during stimulus presentation (bottom). **(B)** Participant-specific classifiers were trained to decode pairs of B categories using multivoxel patterns from the pre-learning run, then applied to A scenes in the post-learning run to test for prediction of the upcoming B category (top). Classifier accuracy per group for hippocampal subfields CA1, CA2/3, and DG (chance = 0.5) (bottom). Dots = individual participants; error bars = bootstrapped 95% CI across participants. \*\**p <* 0.01, \**p <* 0.05

We found successful prediction in both groups. The upcoming B category could be reliably decoded during A images in CA2/3 for both control (*p* = 0.036, 95% CI: [0.50, 0.54]) and stress (*p* = 0.022, 95% CI: [0.50, 0.55]) groups. These effects were specific to CA2/3 and not present in CA1 (control *p* = 0.092, 95% CI: [0.49, 0.53]; stress *p* = 0.188, 95% CI: [0.47, 0.51]) or DG (control *p* = 0.243, 95% CI: [0.47, 0.51]; stress *p* = 0.276, 95% CI: [0.49, 0.52]). As chance classification can deviate from hypothetical levels, we confirmed these results using randomization tests (Supplemental Figure S1). These predictive representations provide offline (i.e., post-learning) neural evidence that both groups were sensitive to statistical regularities, complementing the online evidence from pupillometry.

### 3.4 Acute stress impairs episodic memory for predictive items

Despite comparable indications of statistical learning, we did find differences in episodic encoding for participants exposed to stress. Participants in both groups had above-chance episodic memory for A, B, and X images the next day (A’ vs chance = 0.5; *p <* 0.001; Figure 3A). Examining memory within groups, the control group had significantly better memory for A images (A vs. X: *p* = 0.018, 95% CI = [0.005, 0.052]; B vs. X: *p* = 0.543, 95% CI = [−0.019, 0.035]), a pattern that was not evident in the stress group (A vs. X: *p* = 0.509, 95% CI = [−0.031, 0.016]; B vs. X: *p* = 0.063, 95% CI= [−0.051, 0.001]). Notably, episodic memory was impaired in the stress group relative to the control group for predictive A images (*p* = 0.035, 95% CI: [−0.061, 0.002]), though not for predictable B images (*p* = 0.109, 95% CI: [−0.063, 0.006]) or baseline X images (*p* = 0.752, 95% CI: [−0.024, 0.033]).

**Figure 3:**
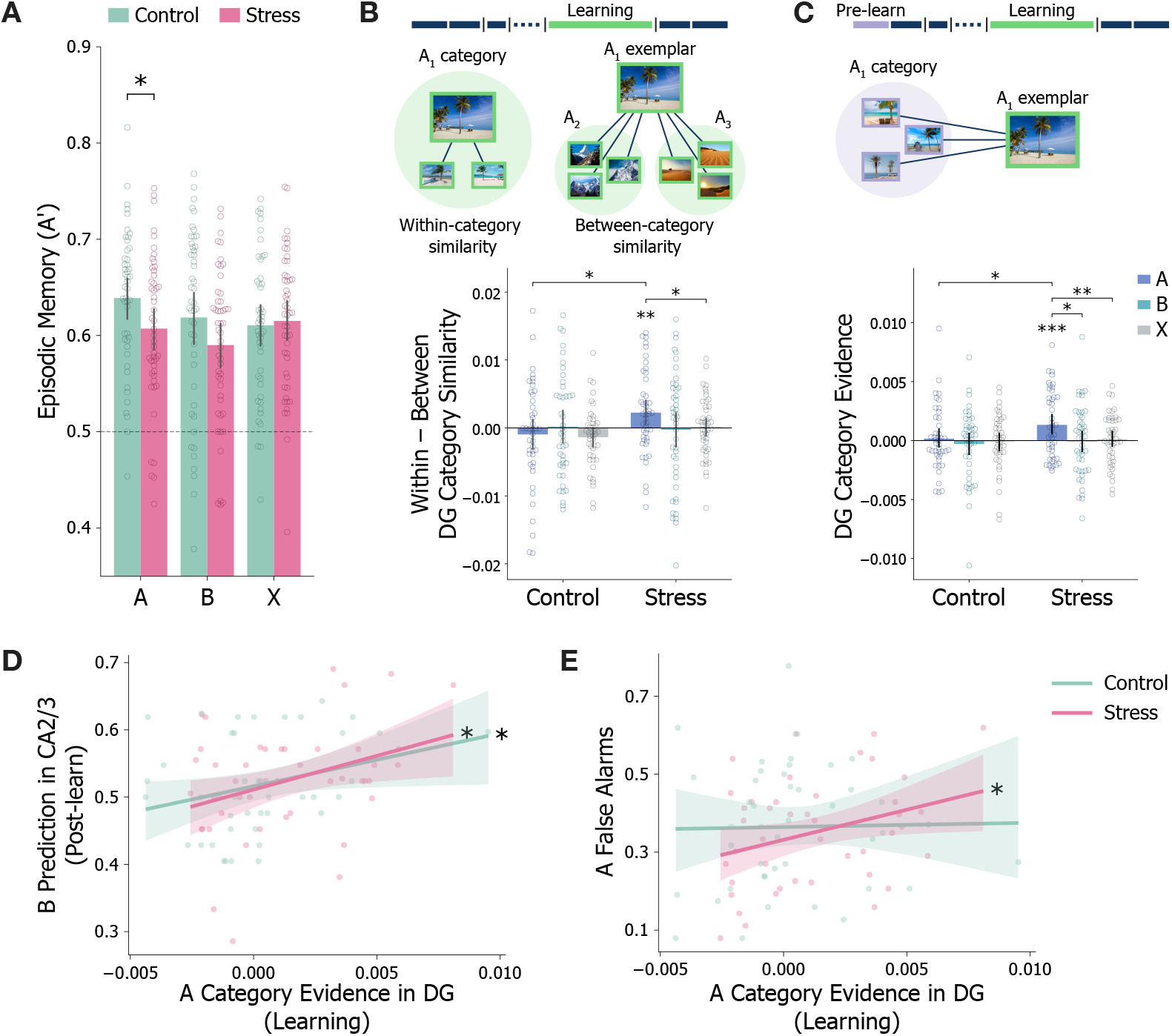
Stress effects on episodic memory. **(A)** Day 2 item recognition memory for A, B, and X scene conditions. **(B)** Pattern separation in dentate gyrus (DG) during learning is assessed by calculating similarity of multivoxel patterns for items within the same vs. different categories from the same condition (top). Difference scores shown per condition and group; values above zero indicate impaired pattern separation (bottom). **(C)** Representation of category information in DG is assessed by calculating multivoxel pattern similarity between each (trial-unique) image during learning with images from the same category in the pre-learning run (top). The resulting value is shown per condition and group, with values above zero indicating category evidence (bottom). **(D)** Relationship between category evidence for A items in DG during learning (see C) and classifier accuracy for the upcoming B item in CA2/3 after learning. **(E)** Relationship between category evidence for A items in DG during learning and false alarm rate for A items. Dots = individual participants; error bars = bootstrapped 95% CI across participants. \*\*\**p <* 0.001, \*\**p <* 0.01, \**p <* 0.05

### 3.5 Acute stress impairs neural pattern separation of predictive items

We examined neural representations during encoding to understand how stress altered episodic memory. Within our task, predictive scenes – which showed impaired episodic memory under stress – may invoke conflicting computations that compete for shared hippocampal resources. To form strong episodic memories, scenes need to be encoded as distinct events [57]. DG supports this process through pattern separation [4, 5]; thus, we would expect DG patterns associated with a scene to be relatively *uncorrelated* with patterns for other scenes from the same category. On the other hand, statistical learning requires processing of commonalities between scenes from the same category to enable extraction of category-level scene transitions. In this case, we would expect patterns associated with a scene to be *more correlated* with other scenes from the same category.

To quantify these processes, we computed mean multivoxel pattern similarity during learning between BOLD activity patterns in DG for scenes from the same category relative to mean pattern similarity with scenes from different categories in the same condition (Figure 3B) [58]. Consistent with pattern separation, this difference was not significantly greater than zero for any condition in the control group (A: *p* = 0.757, 95% CI: [−3.18, 1.47] *×* 10^−3^; B: *p* = 0.475, 95% CI: [−2.18, 2.43] *×* 10^−3^; X: *p* = 0.979, 95% CI: [−2.84, −0.054] *×* 10^−3^), and for B and X conditions in the stress group (B: *p* = 0.583, 95% CI: [−2.81, 2.21] *×* 10^−3^; X: *p* = 0.426, 95% CI: [−1.16, 1.38] *×* 10^−3^).

In contrast, the stress group showed less pattern separation for predictive A scenes, with significantly higher within-than between-category similarity in DG (*p* = 0.006, 95% CI: [0.49, 4.02] × 10^−3^). This result differed significantly from how these same participants processed baseline X scenes (*p* = 0.016, 95% CI: [0.18, 4.12] × 10^−3^) and how the control group processed A scenes (*p* = 0.019, 95% CI: [−0.17, 6.00] × 10^−3^). These effects were not present in hippocampal subfields CA1 or CA2/3 in any condition or group (Supplemental Figure S2).

### 3.6 Hippocampus represents predictive categories under stress

Our results suggest that stress disrupts pattern separation among predictive scenes in DG, which may support processing category-level information useful for predicting the next category. To quantify category-level information directly, we calculated the mean multivoxel pattern similarity between each scene during learning and all scenes from the same category in the pre-learning run, which provides a “category template” unaffected by temporal regularities (Figure 3C).

The findings from this category evidence analysis mirror and extend the pattern separation results. For the control group, category evidence in DG was not significantly greater than zero in any condition (A: *p* = 0.331, 95% CI: [−0.59, 1.04] × 10^−3^; B: *p* = 0.736, 95% CI: [−1.25, 0.59] × 10^−3^; X: *p* = 0.55, 95% CI: [−0.79, 0.66] × 10^−3^). However, the stress group showed significant DG category evidence for predictive A scenes (*p <* 0.001, 95% CI: [0.53, 2.16] × 10^−3^), but not for B (*p* = 0.558, 95% CI: [−0.93, 0.82] × 10^−3^) or X (*p* = 0.354, 95% CI: [−0.50, 0.72] × 10^−3^). Category evidence in the stress group for A was significantly stronger than for B (*p* = 0.016, 95% CI: [0.11, 2.73] × 10^−3^) or X (*p* = 0.006, 95% CI: [0.26, 2.20] × 10^−3^) within participants, as well as category evidence observed in the control group for A (*p* = 0.026, 95% CI: [−2.30, 0.01] × 10^−3^). There was a similar, though marginally significant, pattern of results in CA1 but not in CA2/3 (Supplemental Figure S3).

These results indicate that, although DG does not typically represent scene categories, stress drives DG representations to code for this information, perhaps because it is relevant to prediction. To test whether category evidence is indeed beneficial to statistical learning, we examined whether A category evidence in DG during learning was associated with subsequent prediction of B in CA2/3 after learning (Figure 2D). Indeed, stronger A category evidence positively correlated with prediction in both groups (stress: *r*(43) = 0.35, *p* = 0.024; control: *r*(40) = 0.29, *p* = 0.038). On the other hand, representing A scenes at a categorical level could impair encoding of idiosyncratic features necessary for episodic memory. Consistent with this idea, stronger A category evidence tracked more false alarms in the stress group (*r*(43) = 0.32, *p* = 0.040; Figure 3E) but not the control group (*r*(40) = 0.019, *p* = 0.907). Together, these results suggest that stress leads the hippocampus to represent statistically predictive information at the expense of episodic memory precision.

### 3.7 Stress prioritizes statistical prediction in monosynaptic pathway

Beyond individual subfields, hippocampal circuit pathways arbitrate between statistical learning and episodic encoding. In particular, the direct connection between CA1-EC is common to both TSP (episodic encoding) and MSP (statistical learning); thus, activity along this pathway could potentially reflect a trade-off between these memory processes. If CA1-EC supports episodic encoding, we would expect differential cofluctuation for scenes that were subsequently remembered vs. forgotten. On the other hand, if CA1-EC supports statistical learning, we would expect differential cofluctuation for scenes in pairs that were subsequently retained in the pair familiarity test. Here we test how stress influences the representation of statistical and episodic encoding within this overlapping pathway.

As stress primarily impacted hippocampal processing of predictive A items, we used an edge timeseries approach to isolate momentary cofluctuation between CA1 and EC during A scenes. Momentary cofluctuation in CA1-EC was related to both episodic encoding and statistical learning, but these relationships were modulated by stress (Figure 4). In the control group, CA1-EC cofluctuation tracked episodic encoding, with higher cofluctuation during subsequently remembered compared to forgotten A items (Figure 4A; *F* (1, 41) = 5.33, *p* = 0.026). However, this was not evident in the stress group (*F* (1, 44) = 2.16, *p* = 0.148; subsequent memory *×* group: *F* (1, 85) = 7.11, *p* = 0.009).

**Figure 4:**
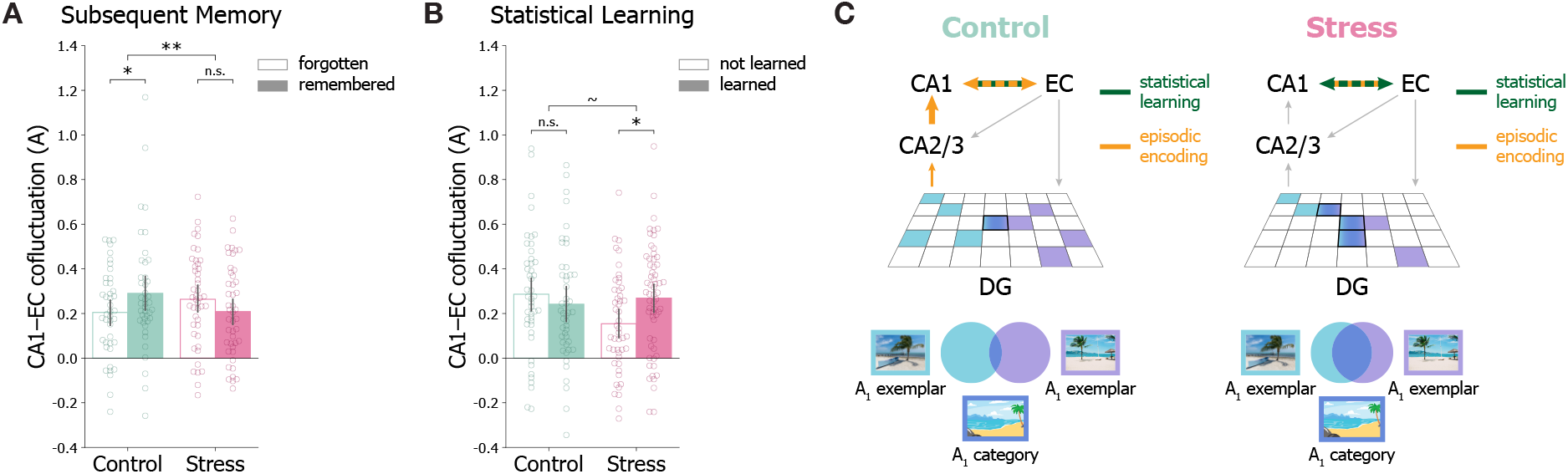
Stress effects on processing in the monosynaptic pathway. **(A)** Momentary cofluctuation between CA1 and EC for predictive A items that were subsequently remembered and forgotten within each group. Dots = individual participants. **(B)** Momentary cofluctuation between CA1 and EC for predictive A items belonging to pairs that are either learned (above-chance pair familiarity strength) or not learned (below-chance) within each group. Dots = individual A-B pairs (3 per participant). **(C)** Schematic summarizing the effects of stress on statistical (green) and episodic (yellow) processing of predictive scenes in the hippocampus. In the control group, connectivity between CA1-EC, as well as TSP connections DG-CA2/3 and CA2/3-CA1, relates to episodic memory but not statistical learning (see Supplemental Figure S5). Under stress, connectivity between CA1-EC relates to statistical learning but not episodic memory, and DG representations show increased (predictive) category information. Error bars = 95% CI across participants. \*\**p <* 0.01, \**p <* 0.05, ∼*p <* 0.07

Instead, CA1-EC cofluctuation in the stress group tracked statistical learning. For the purposes of visualization and comparison with episodic encoding, we binarized performance on the pair familiarity test (above-chance accuracy = “learned”; below-chance accuracy = “not learned”). During A items, CA1-EC cofluctuation in the stress group was significantly higher for learned compared to not learned pairs (Figure 4B; *F* (1, 62) = 4.75, *p* = 0.033). This pattern was not evident in the control group (*F* (1, 49) = 0.51, *p* = 0.480; familiarity × group: (*F* (1, 111) = 3.69, *p* = 0.057). A version of this analysis using a continuous measure of pair familiarity strength revealed an even stronger relationship between CA1-EC and statistical learning with stress (Supplemental Figure S4A), which was also evident at the participant level (Supplemental Figure S4B).

Consistent with its shared role in both TSP and MSP, these results show that CA1-EC cofluctuation reflects a trade-off between episodic memory and statistical learning. Accordingly, we expect connections exclusive to TSP to support episodic encoding, but not statistical learning. Indeed, neither group shows a positive association between cofluctuation and statistical learning along any TSP connection – namely, EC-DG, EC-CA2/3, DG-CA2/3, and CA2/3-CA1 (Supplemental Figure S5B). In fact, stronger cofluctuation between CA2/3 and CA1 is negatively associated with statistical learning in the control group. In contrast, cofluctuation in CA2/3-CA1, as well as DG-CA2/3, is positively associated with episodic memory in the control group, but not the stress group (Supplemental Figure S5A). Together, these results complement earlier subfield analyses with a circuit-based mechanism by which stress prioritizes processing of statistical regularities at the expense of episodic details (Figure 4C).

## 4 Discussion

Here we show how stress, known to impair episodic memory, influences hippocampal processes associated with statistical learning. We find that acute stress impairs episodic memory for predictive items, reducing hippocampal processes like pattern separation in favour of representing features beneficial for prediction. We further demonstrate that stress alters hippocampal circuit dynamics, biasing CA1-EC connectivity away from episodic encoding towards statistical learning. Together, these findings uncover new learning benefits under stress, with attenuated episodic encoding reflecting a shift toward statistical prediction.

Our findings complement work in non-human animals showing hippocampal subfield-specific effects of stress and glucocorticoids, and extend these findings by showing their association with human learning. Namely, we identified two hippocampal mechanisms by which stress prioritized predictive processing over episodic encoding: reduced pattern separation in DG, and strengthened relationships between CA1-EC coupling and learning predictive regularities. We consider each of these mechanisms below.

In the stress group, representations of predictive scenes in DG became less distinctive and more categorical, capturing information that helps predict the upcoming scene. This increased category evidence was associated with stronger hippocampal prediction and weaker episodic memory precision. These findings are consistent with past work showing that acute stress shifts memory processing towards gist-like information and away from fine-grained details [59, 60, 61], a bias associated with altered DG engram cells in a rodent model [62]. Our task design highlights a situation in which a shift toward gist-level processing may be particularly adaptive. Namely, representing predictive stimuli in a gist-like manner (i.e., with categorical features rather than idiosyncratic details) facilitates learning which scene category will occur next. We note that, without stress, episodic memory for predictive items can be retained even when statistical regularities are learned (consistent with [20]). Although a shift toward gist-level encoding is commonly characterized as a transition from hippocampal to cortical systems [63, 59], our results reveal that a similar shift can occur across subsystems within the hippocampus [64]. They also suggest that stress may confer benefits in other situations in which identifying links between experiences comes at the expense of remembering individual event details [65].

In addition to altering DG representations, stress changed the role of CA1-EC connectivity. We propose that this pathway, common to both MSP (associated with statistical learning) and TSP (associated with episodic encoding), serves as a locus of competition between these two forms of learning. By taking a time-locked approach and calculating CA1-EC coupling at predictive moments, we showed that the CA1-EC pathway can flexibly support either episodic or statistical learning. Under stress, weakened pattern separation in DG may lead to more prominent category representation for predictive items spreading through TSP and eventually through CA1-EC. This may further reinforce category representation in DG, as neural network models show that category representation can spread through EC back to DG through big-loop recurrence [66, 67]. These findings underscore the need to consider modulatory factors like stress in network models of hippocampal function.

Beyond stress, evaluating hippocampal subfields also allows for precise localization of hippocampal contributions to encoding [68, 69]. We found predictive representations of upcoming B categories during A in CA2/3 only, as well as stress-induced category evidence for A only in DG. The lack of prediction or category evidence in CA1 is surprising at first glance, given its role in statistical learning as a key component of MSP. However, a recent account proposes that prediction is facilitated by pattern completion, enabling retrieval of associated memory representations based on partial information [70]. This function has been attributed to CA3, which contains unique recurrent connections forming a highly interconnected autoassociative network [71, 72, 73]. Our results contribute to a growing body of experimental evidence in support of this account, including recent findings in 7T fMRI identifying representations of visual predictions only in CA2/3 [56], as well as related evidence of prediction in CA2/3/DG from studies that did not separate these subfields [55, 2]. Furthermore, we found that participants in the stress group showed only marginal evidence of category representation in CA1 with a much stronger effect in DG. This deviates from category-learning paradigms, with evidence from both neuroimaging and neural network modelling suggesting that CA1 has more similar representations for items within the same category [74, 66]. This discrepancy may be related to differences in task demands. For example, CA1 has been theorized to support comparison between predicted and experienced inputs [75], and may support non-directional associative representations [2, 76] and prediction error signals [77]. Thus, although CA1 may represent category-level content when initially forming category knowledge, it may instead support such predictive comparisons in tasks like the present one that involve temporal predictions. Further work is needed to clarify the varied roles of CA1 in learning and representing predictive regularities and how these processes may be modulated by stress.

In addition to successful hippocampal prediction of upcoming scenes, we found that pupil responses during learning reflected sensitivity to predictive regularities. In both groups, pupil size differentiated predictive A and predictable B images. Prediction has well-documented effects on pupil size: unexpected or prediction-violating stimuli in both auditory [78] and visual modalities [79] evoke increased pupil dilation (or less constriction) compared to predicted stimuli, though effects are weaker in the visual modality [80]. Pupil size also plays a role in detecting temporal structure [81] and regularities [82]; however, to our knowledge prediction has not previously been measured using pupillary response during statistical learning. As the stimuli in our study were visual, the dominant pupil response is one of constriction due to the change in luminance. Thus, we would expect more constriction for predicted stimuli, and indeed observed this pattern in the control group. However, although the stress group also differentiated predictive and predictable images, the pattern was reversed, with stronger constriction for predictive A items. Given the roles of pupil responses in arousal [83], and interactions between pupil-associated adrenergic responses and other parts of the stress response [84, 85, 86, 85], it is reasonable to expect that stress might modulate pupil responses during learning. Previous work suggests that stress exposure can lead to more stimulus-evoked pupil constriction [87] and, in threatening situations, that more constriction reflects more certainty [88]. Furthermore, recent work suggests that pupil dilation tracks interactive effects of salience and predictability, with less constriction for predictable elements of salient musical sequences [89]; this may be related to less constriction for predictable B items under stress. Further work is required to disentangle how stress modulates pupil response to learning predictive regularities.

Our findings reveal an adaptive role for stress in changing hippocampal mechanisms to prioritize predictive regularities. By suppressing encoding of idiosyncratic details and dedicating neural resources towards predictive information, stress may enhance our ability to respond to unexpected events. This study also adds to the growing literature demonstrating that multiple learning functions are supported by competing hippocampal pathways. We show that these pathways can be differentially modulated by stress, highlighting a dynamic that may uncover opportunities for enhancing learning under stress.

## Data and code availability

Data and code are available by request, and will be released publicly upon acceptance.

## Author contributions

I.Z.: conceptualization, data curation, formal analysis, investigation, methodology, software, validation, visualization, writing–original draft, writing–review and editing; Z.B.: data curation, investigation, software, writing–review and editing; Y.H.: investigation, methodology, writing–review and editing; E.G.W.: investigation, writing–review and editing; B.E.S.: conceptualization, methodology, resources, writing–review and editing; N.B.T.-B.: conceptualization, supervision, writing–review and editing; E.V.G.: conceptualization, funding acquisition, investigation, methodology, project administration, resources, supervision, writing–original draft, writing–review and editing.

## Supplementary Figures

**Figure S1:**
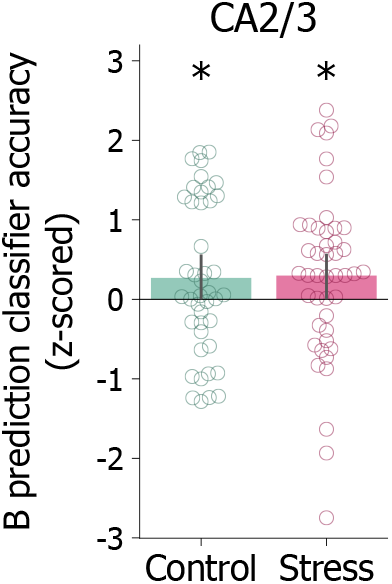
Predictive category decoding analysis with randomization testing. Z-scored classifier accuracies for hippocampal subfield CA2/3 are displayed for each group (see also main text Fig. 2B). Classifier accuracies are z-scored relative to participant-specific null distributions of accuracy values. Z-scored classifier accuracies are reliably above zero for participants in both control (*p* = 0.033, 95% CI: [−0.018, 0.559]) and stress (*p* = 0.027, 95% CI: [−0.005, 0.597]) groups. Dots = z-scored classification accuracies from individual participants. Error bars = bootstrapped 95% CIs across participants. \**p <* 0.05

**Figure S2:**
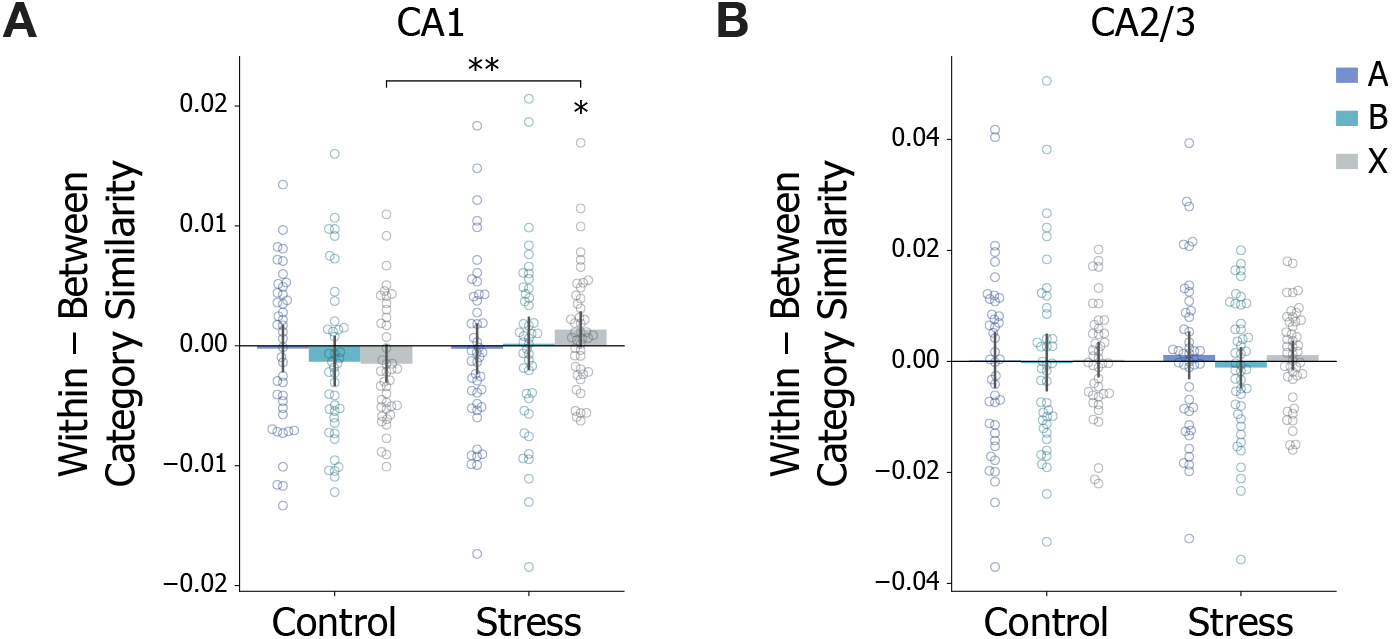
Category representation similarity analysis applied to hippocampal subfields CA1 and CA2/3. DG results shown in main text Fig. 3B. The difference between within- and between-group pattern similarity during learning is shown for each condition and group. **(A)** In CA1, this difference is not significantly greater than zero for any condition in the control group (A: *p* = 0.591, 95% CI: [− 2.15, 1.67] × 10^−3^; B: *p* = 0.903, 95$ CI: [−3.29, 0.69] × 10^−3^; X: *p* = 0.972, 95% CI: [*−*2.97, 0.03] × 10^−3^), and for the A and B conditions in the stress group (A: *p* = 0.597, 95% CI: [−2.26, 1.82] *×* 10^−3^; B: *p* = 0.437, 95$ CI: [−1.94, 2.33] *×* 10^−3^). However, the difference is significantly higher for the X condition in the stress group (*p* = 0.025, 95% CI: [0.00, 2.77] *×* 10^−3^), compared with the X condition in the control group (*p* = 0.003, 95% CI: [0.81, 4.89] *×* 10^−3^). **(B)** In CA2/3, this difference is not significantly greater than zero for any condition in the control (A: *p* = 0.476, 95% CI: [−4.62, 5.05] *×* 10^−3^; B: *p* = 0.557, 95% CI: [−5.07, 4.77] *×* 10^−3^; X: *p* = 0.424, 95% CI: [−2.65, 3.20] *×* 10^−3^) or stress (A: *p* = 0.293, 95% CI: [−2.92, 5.31] *×* 10^−3^; B: *p* = 0.733, 95$ CI: [−4.68, 2.28] *×* 10^−3^; X: *p* = 0.182, 95% CI: [−1.32, 3.56] *×* 10^−3^) groups. Dots = individual participants. Error bars = bootstrapped 95% CIs. \*\**p <* 0.01, \**p <* 0.05

**Figure S3:**
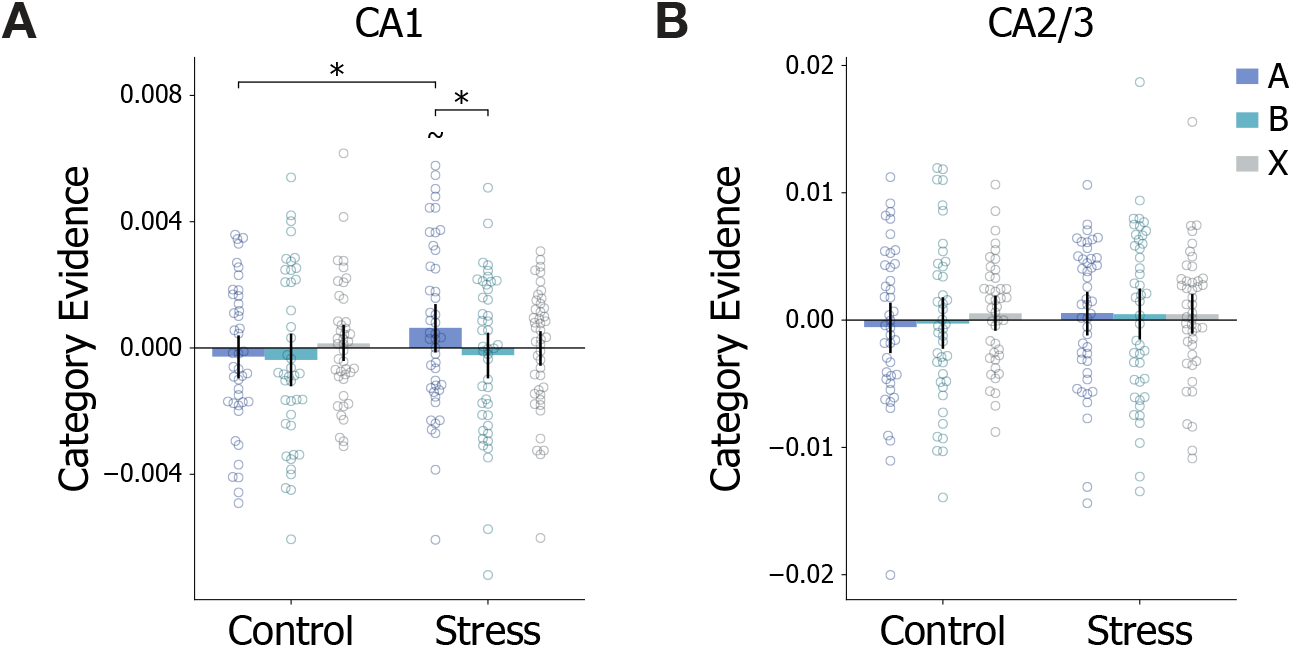
Category evidence analysis applied to hippocampal subfields CA1 and CA2/3, with category evidence shown for each condition and group. **(A)** Category evidence in CA1 was not significantly greater than zero in any condition in the control group (A: *p* = 0.778, 95% CI: [*−*0.97, 0.41] *×* 10^−3^; B: *p* = 0.815, 95$ CI: [*−*1.20, 0.46] *×* 10^−3^; X: *p* = 0.317, 95% CI: [*−* 0.41, 0.74] *×* 10^−3^). The stress group showed marginally significant CA1 category evidence for predictive A scenes (*p* = 0.054, 95% CI: [−0.14, 1.42] × 10^−3^), but not for B (*p* = 0.730, 95% CI: [*−*0.96, 0.48] × 10^−3^) or X (*p* = 0.494, 95% CI: [*−*0.57, 0.54] × 10^−3^). CA1 category evidence for A was significantly stronger than for B (*p* = 0.041, 95% CI: [*−*0.10, 1.87] × 10^−3^) but not X (*p* = 0.101, 95% CI: [*−*0.34, 1.63] × 10^−3^) within the stress group, and was also significantly stronger than category evidence for A in the control group (*p* = 0.043, 95% CI: [*−*0.13, 1.95] × 10^−3^). This pattern of results is similar to (but weaker than) those found in DG (see main text Fig. 3C), and suggest that CA1 may also partially represent the category of predictive items under stress. **(B)** Category evidence in CA2/3 was not significantly greater than zero in any condition for either the control (A: *p* = 0.715, 95% CI: [−2.48, 1.32] *×* 10^−3^; B: *p* = 0.611, 95% CI: [−2.20, 1.67] *×* 10^−3^; X: *p* = 0.206, 95% CI: [−0.74, 1.81] *×* 10^−3^) or stress (A: *p* = 0.243, 95% CI: [−1.08, 2.12] *×* 10^−3^; B: *p* = 0.322, 95% CI: [−1.47, 2.38] *×* 10^−3^; X: *p* = 0.265, 95% CI: [−0.99, 1.92] *×* 10^−3^) groups. Dots = individual participants. Error bars = bootstrapped 95% CIs. \**p <* 0.05, ∼*p <* 0.07

**Figure S4:**
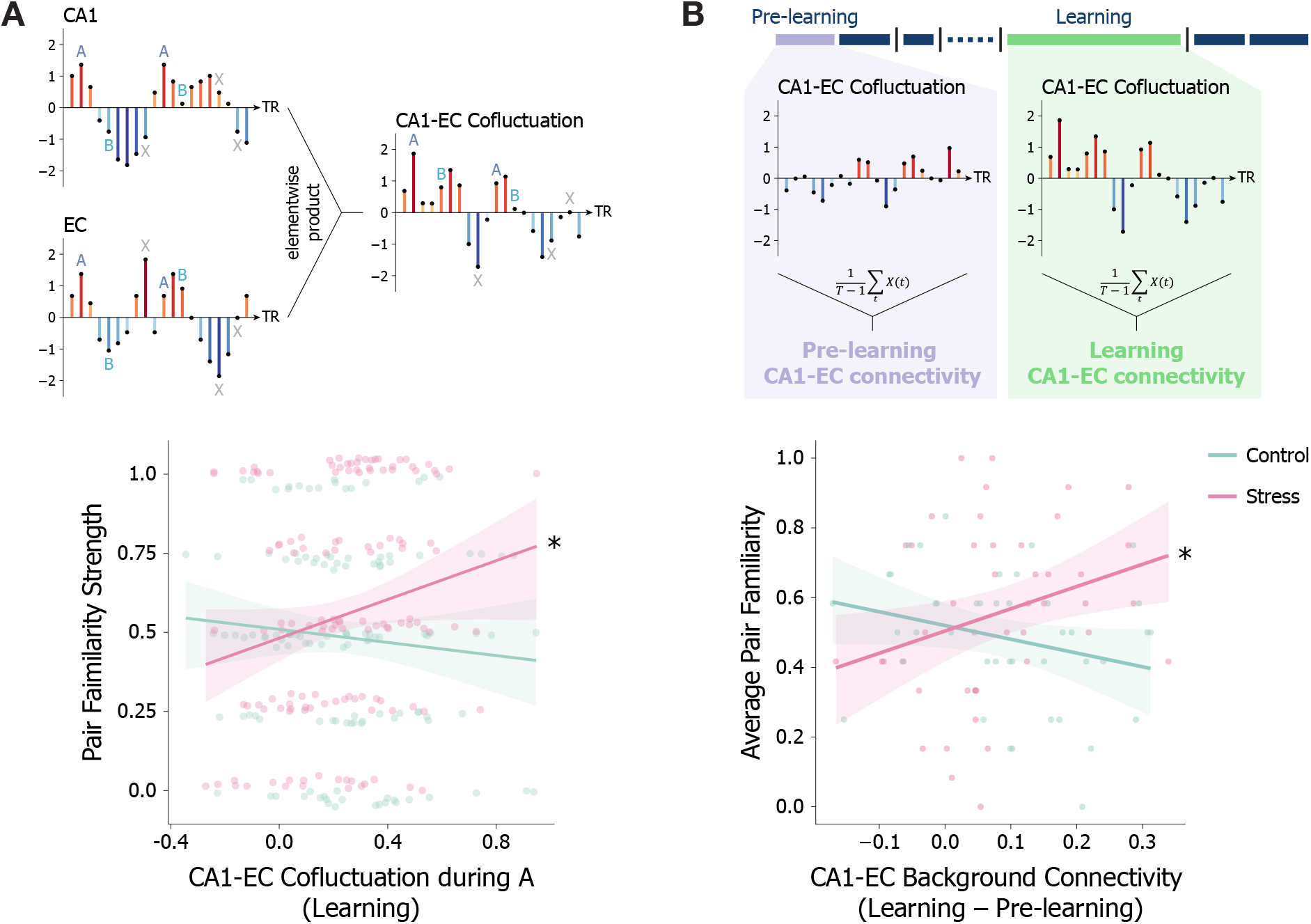
Connectivity analyses with pair familiarity strength as a continuous variable. **(A)** Momentary cofluctuation between CA1-EC is computed by taking the elementwise product of the z-scored mean timeseries from CA1 and EC during learning. Timepoints that occur during A images (offset by 3 TRs to account for haemodynamic lag) are used for analyses (top). The relationship between mean CA1-EC cofluctuation during each A category and pair familiarity strength for the corresponding category pair is shown for each group. Stronger CA1-EC cofluctuation was positively associated with pair familiarity strength in the stress group (*F* (1, 89) = 4.95, *p* = 0.029), but not in the control group (*F* (1, 83) = 1.30, *p* = 0.26; cofluctuation × group interaction (*F* (1, 172) = 5.86, *p* = 0.017). Dots = individual A-B pairs (3 per participant). Solid lines = predicted values from within-group LMEs. Error shading = bootstrapped 95% CIs over LME prediction values from 9,999 iterations of within-group participant resampling. Binary version of this analysis shown in main text Fig. 4B. **(B)** We employed a background connectivity approach to examine whether overall CA1-EC connectivity during learning is related to statistical learning at the participant level. As with the momentary cofluctuation analysis, we first used a GLM to regress out stimulus-evoked responses. We then extracted the mean of the model residuals across voxels in CA1 and EC. Background connectivity was finally calculated as the Pearson correlation of these mean time-series, normalized using a Fisher r-to-z transform. For each run, this value is equivalent to the mean of momentary cofluctuations across time (more specifically, the sum of the cofluctuation timeseries *X*(*t*) divided by the total number of timepoints *T* minus 1) (top). The difference between CA1-EC background connectivity during the learning and pre-learning runs is positively associated with average pair familiarity performance in the stress group (*r*(43) = 0.26, *p* = 0.023), but not in the control group (*r*(40) = *−*0.26, *p* = 0.068). Pre-learning connectivity is subtracted from learning connectivity to isolate changes in connectivity elicited by statistical learning (bottom). Dots = individual participants. Error shading = bootstrapped 95% CIs. \**p <* 0.05

**Figure S5:**
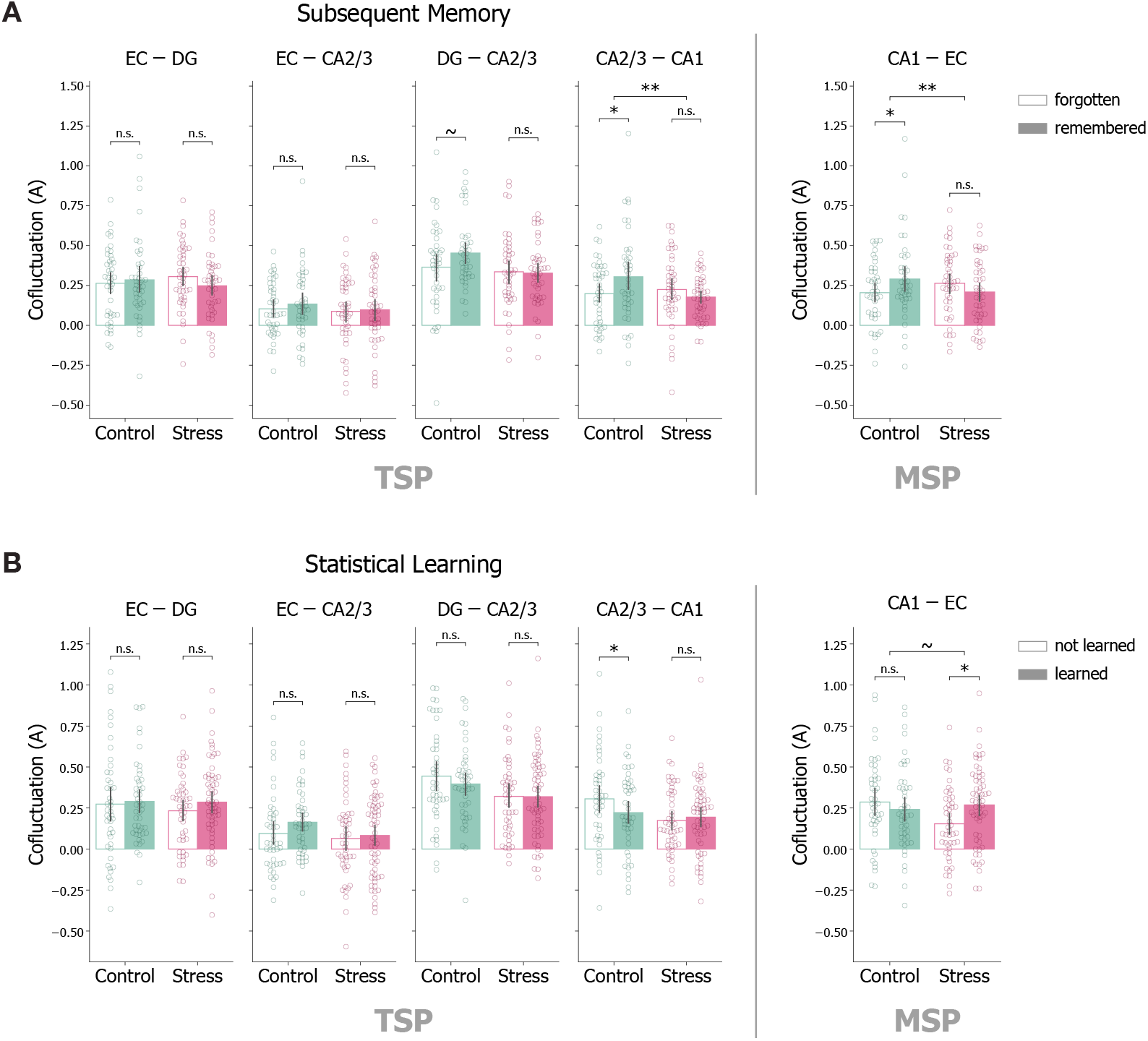
Momentary cofluctuation analyses along trisynaptic pathway connections. **(A)** Momentary cofluctuation between pairs of ROIs corresponding to TSP connections (EC-DG, EC-CA2/3, DG-CA2/3, CA2/3-CA1) for predictive A items that were subsequently remembered and forgotten within each group. For EC-DG and EC-CA2/3, cofluctuation was not significantly related to episodic memory in either the control (EC-DG: *F* (1, 41) = 0.25, *p* = 0.617; EC-CA2/3: *F* (1, 41) = 0.53, *p* = 0.471) or stress (EC-DG: *F* (1, 44) = 1.96, *p* = 0.168; EC-CA2/3: *F* (1, 44) = 0.056, *p* = 0.813) groups. DG-CA2/3 cofluctuation was marginally related to episodic memory in the control group, with higher cofluctuation for subsequently remembered compared to forgotten images (*F* (1, 41) = 3.56, *p* = 0.066), but there was no apparent association between DG-CA2/3 cofluctuation and episodic memory in the stress group (*F* (1, 44) = 0.034, *p* = 0.855; subsequent memory × group: *F* (1, 85) = 2.19, *p* = 0.142; main effect of group: *F* (1, 85) = 4.12, *p* = 0.045). CA2/3-CA1 cofluctuation was also associated with episodic memory in the control group, with significantly higher cofluctuation for subsequently remembered compared to forgotten images (*F* (1, 41) = 6.47, *p* = 0.015), but this relationship was again not evident in the stress group (*F* (1, 44) = 1.62, *p* = 0.210; subsequent memory × group: *F* (1, 85) = 7.56, *p* = 0.007). Given the positive associations between TSP cofluctuation (in particular, along TSP connections downstream of DG) and episodic memory, the lack of effect in the stress group suggests that stress may impair episodic memory processing along TSP. For comparison, CA1-EC results are shown on the right. Dots = individual participants. **(B)** Momentary cofluctuation along TSP connections for predictive A items belonging to pairs that are either learned or not learned within each group. For EC-DG, EC-CA2/3, and DG-CA2/3, cofluctuation was not significantly related to statistical learning in either the control (EC-DG: *F* (1, 49) = 0.013, *p* = 0.909; EC-CA2/3: *F* (1, 49) = 2.25, *p* = 0.140; DG-CA2/3: *F* (1, 49) = 0.90, *p* = 0.346) or stress (EC-DG: *F* (1, 62) = 1.16, *p* = 0.286; EC-CA2/3: *F* (1, 62) = 0.004, *p* = 0.952; DG-CA2/3: *F* (1, 62) = 0.001, *p* = 0.982) groups. CA2/3-CA1 cofluctuation is negatively associated with statistical learning in the control group, with lower cofluctuation for images belonging to learned pairs (*F* (1, 49) = 4.44, *p* = 0.040), while no relationship was evident in the stress group (*F* (1, 62) = 0.23, *p* = 0.631; familiarity × group: *F* (1, 111) = 3.10, *p* = 0.081; main effect of group: *F* (1, 85) = 4.32, *p* = 0.041). The lack of positive associations between TSP cofluctuation and statistical learning is consistent with the hypothesized role of TSP. For comparison, CA1-EC results are shown on the right. Dots = individual A-B pairs (3 per participant). Error bars = bootstrapped 95% CIs. \*\**p <* 0.01, \**p <* 0.05, ∼*p <* 0.07

